# Proteomic Characterization of Striatal Neurabin Interactome and its Sex Specific Impact on Motor Behavior

**DOI:** 10.1101/2025.10.09.681445

**Authors:** Nikhil R. Shah, Wesley B. Corey, Camden N. Harris, Anthony J. Baucum

**Affiliations:** Medical Neuroscience Graduate Program, Indiana University School of Medicine; Department of Biology, Indiana University Indianapolis; Neuroscience Experience and Undergraduate Research Opportunities Program, Indiana University Indianapolis; Department of Biochemistry, Molecular Biology, and Pharmacology, Indiana University School of Medicine; Stark Neurosciences Research Institute, Indiana University School of Medicine

## Abstract

The striatum serves as the primary input nucleus of the basal ganglia. Reversible protein phosphorylation in the post synaptic density (PSD) of medium spiny neurons (MSNs) modulates inputs from striatal afferents. The context dependent regulation of PSD protein phosphorylation in direct-pathway medium spiny neurons (dMSNs) and indirect-pathway medium spiny neurons (iMSNs) works to differentially and synergistically impact striatal physiology and the execution of motor programs. An important regulator of PSD protein phosphorylation is protein phosphatase 1 (PP1), which obtains substrate specificity through the action of PP1 targeting proteins. While prior work has demonstrated the global and cell type-specific impact of the PP1 targeting protein, spinophilin, on striatal motor behaviors like the accelerating rotarod task and amphetamine sensitization, the role of its homologue, neurabin, is yet to be elucidated. Proteomics determined that striatal neurabin associates with pre and postsynaptic proteins associated with glutamatergic synapse function. Moreover, we found that global loss of neurabin enhanced rotarod motor learning but had no impact on amphetamine sensitization. Interestingly, using novel conditional neurabin knockout mouse lines, we found that loss of neurabin in dMSNs, but not iMSNs, enhanced performance on the accelerating rotarod task and that these effects were specific for male mice. These data highlight neurabin’s particular importance to the striatal glutamatergic synapse and uncover a sex and cell type specific role for this synaptic protein in uniquely limiting skill motor learning but not psychomotor sensitization.

## Introduction

The basal ganglia consist of several subcortical nuclei with roles in motor learning, decision making, and reward processing^1^. Anatomically, the striatum is the largest component of the basal ganglia and serves as its primary input nucleus^1,2^. A central function of the striatum is to integrate information from myriad brain regions to facilitate motor learning and motor output^1–3^. This process largely occurs through the action of striatal medium spiny neurons (MSNs), a prominent class of GABAergic projection neurons, and one of the striatum’s primary cell types^4^. Direct and indirect pathway medium spiny neurons (dMSNs and iMSNs, respectively) serve as a crucial junction between striatal afferents and the remainder of the basal ganglia^5–7^.

Reversible protein phosphorylation within the post-synaptic density (PSD) is an important component of processing afferent signaling. The phosphorylation states of PSD proteins are regulated by a suite of kinases and phosphatases^8–10^. While protein kinases are varied and largely substrate specific^11^, protein phosphatases are relatively homogenous^12^ and require the role of targeting proteins to obtain substrate specificity^13^. Spinophilin and neurabin are the two most abundant PSD-enriched actin binding and protein phosphatase 1 (PP1) targeting proteins^14^. PP1 is a prominent protein phosphatase that regulates neuronal signaling^15^. Spinophilin and neurabin are highly homologous, and both co-localize and co-immunoprecipitate^16–18^. Despite their homology and interaction, studies have delineated distinct, yet complementary, roles for spinophilin and neurabin in regulating cell signaling, spine density, and rodent fear and anxiety behaviors^19–23^. In the striatum, spinophilin and neurabin enforce opposing forms of corticostriatal plasticity and may modulate drug reward by differentially impacting behavioral and biochemical sensitization to cocaine^24^. Furthermore, studies characterizing the spinophilin-specific striatal interactome have delineated the global and cell type-specific impact of spinophilin on striatal associated motor behaviors including amphetamine sensitization, rotarod motor learning, and compulsive grooming^20,25,26^. Despite neurabin’s established roles in striatal physiology^24,27,28^, and its putative importance in regulating striatal output, neurabin’s striatal interactome has yet to be characterized and its cell type-specific impact on striatal associated behaviors is unknown. Using our recently generated global and conditional neurabin knockout mice, we define neurabin’s striatal interactome and determine that global loss of neurabin has no effect on amphetamine sensitization, but that global and dMSN-cell type specific loss of neurabin enhances rotarod motor learning, uncovering neurabin as a brake on motor performance, but not amphetamine sensitization.

## Methods

### General Animal Maintenance

Mice were maintained with ad libitum food and water on a 7AM-7PM light dark cycle. All animal rotarod studies were performed between 11 AM and 6 PM. All studies were performed in accordance with Indiana University Indianapolis School of Science or School of Medicine Institutional Animal Care and Use Committees (Protocol #s SC310 (School of Science) and 21090, 22024, and 25030 (IU School of Medicine)).

### Generation of Global Neurabin and Conditional Neurabin KO Mice

Mice bearing loxp sites around exon 6 of the neurabin locus (Nrb^fl/fl^) were generated by the University of Michigan Transgenic Animal Model Core using Easi CRISPR-mediated homology-dependent repair of the neurabin gene, *Ppp1r9a*, in mouse embryonic stem cells (ES) as previously described ^29^. Non-homologous end joining using these approaches resulted in a frame shift mutation around exon 6 and the generation of a null neurabin allele that provided us with a neurabin global knockout line (Nrb^−/−^) ^30^. Conditional knockout mice were validated by crossing Nrb^fl/fl^ mice with CagCreER (Tg(CAGcre/Esr1*)5Amc Jackson Laboratories #004682), a line carrying a global, tamoxifen-inducible Cre recombinase to generate Nrb^CagCreER^ mice. We also obtained Adora2a-Cre (MMRRC strain 036158-UCD, B6.FVB(Cg)-Tg(Adora2a-cre)KG139Gsat/Mmucd) and D1 Cre (036916-UCD B6.FVB(Cg)-Tg(Drd1acre)FK150Gsat/Mmucd) from the MMRRC. We crossed mice hemizygous for D1-Cre (Cre^D1^), which is Cre expressing only in cells with D1 receptors, and Adora2a-Cre (Cre^A2a^), which is Cre expressing only in cells with adenosine A2a receptors, with Nrb^fl/fl^ mice to obtain Nrb^ΔdMSN^ and Nrb^ΔiMSN^ mice, respectively. Primer sequences for genotyping can be found in **Table S1**.

### Neurabin immunoprecipitation, tryptic digestion, and mass spectrometry

Two adult neurabin wildtype (WT) and two global knockout (KO) male mice were used for these studies. Striata were dissected from these mice, homogenized in a dounce homogenizer, sonicated, and then homogenates centrifuged at 13,600 x g at 4ºC. For immunoprecipitation (I.P.), 500 µL of each lysate was transferred to a clean microfuge tube and 2 µg of polyclonal rabbit anti-neurabin antibody (Santa Cruz Biotechnology Inc, Dallas, TX, sc-32932; discontinued) was added to each lysate. Samples were rocked overnight at 4°C. The following morning, 20 µL of Protein G Dynabeads (Thermo-Fisher Scientific, Waltham, MA, 10003D) were added to each IP and samples were rocked for 2 hours at 4°C. After rocking, the samples were washed with IP wash buffer (50mM Tris HCL, 150 mM NaCl, 0.5% Triton X-100) 3 times and rocked for 5 minutes at 4°C in between each wash. After the wash steps, the beads were suspended in 40 µL of PBS. Thirty µL of 8 M Urea in 100 mM Tris pH 8.5 was added to each sample and proteins were reduced with 5 mM tris(2-carboxyethyl)phosphine hydrochloride (TCEP, Sigma-Aldrich, St. Louis, MA) for 30 minutes at room temperature. The resulting free cysteine thiols were alkylated with 10 mM chloroacetamide (CAA, Sigma Aldrich) for 30 min at room temperature in the dark. Samples were diluted with 50 mM Tris-HCl, pH 8.5 to a final urea concentration of 2 M. Samples were then digested overnight at 35ºC with 0.5 µg Trypsin/Lys-C (Mass Spectrometry grade, Promega Corporation). Digestions were quenched and acidified with formic acid (FA, 1% v/v; Honeywell, Charlotte, NC).

Fifteen µl of each sample was desalted on a 5 cm C18 trap column Acclaim™ PepMap™ 100 (3 µm particle size, 75 µm diameter; Thermo Fisher Scientific) followed by separation on a 15 cm PepMap RSLC C18 EASY-Spray column (Thermo Fisher Scientific) using an UltiMate 3000 HPLC coupled to a Q-Exactive Plus mass spectrometer (Thermo Fisher Scientific) operated in positive ion mode. The gradient used for the separation of the peptides was 5-35% solvent B over 60 mins, 35-80% solvent B over 2 mins, 80-95% solvent B over 6 mins, and 95-3% solvent B for 2 mins (Solvent A: 95% water, 5% acetonitrile, 0.1% FA; Solvent B: 100% acetonitrile, 0.1% FA). A data-dependent TopN20 mass acquisition method was used with the following parameters for the MS1: mass resolution: 70,000; AGC target: 3e6; maximum IT: 100 ms; and mass scan range: 300-2000 m/z. MS2 parameters were fixed first mass: 100 m/z; resolution: 17,500; AGC target: 1e5; maximum IT: 50 ms; isolation window: 4.0 m/z; normalized collision energy: 30; dynamic exclusion: 30 s; and charge exclusion: 1, 7, 8, >8.

### Proteomics analysis of mass spectrometry

Raw files were analyzed in Proteome Discover™ 2.5 (Thermo Fisher Scientific) with a database containing Uniprot reviewed and unreviewed Mus musculus as well as common contaminants (Total sequences: 49912). Sequest HT searches were conducted with full trypsin, a maximum of 3 missed cleavages, precursor mass tolerance of 10 ppm, and fragment mass tolerance of 0.02 Da. Static modifications used for the search were carbamidomethylation on cysteine (C) residues; Dynamic modifications used for the search were oxidation of methionines, deamidation of asparagine, and phosphorylation on serine/threonine/tyrosine. Dynamic Protein terminus modifications allowed were acetylation (N-terminus), Met-loss, or Met-loss plus acetylation (N terminus). Percolator False Discovery Rate was set to a strict setting of 0.01 and a relaxed setting of 0.05, and the Protein FDR validator in the consensus was set to a strict 1% protein FDR cutoff and relaxed 5% protein FDR cutoff. Search results were loaded into Scaffold (version Scaffold_5.0.1, Proteome Software Inc., Portland, OR) with prefiltered FDR (percolator output) for viewing and statistical analysis. Data were filtered for proteins containing at least two unique peptides. A t-test based on total spectral counts with Benjamini-Hochberg test correction was performed to compare KO and WT IP samples. 1005 neurabin striatal interacting proteins (NrbAPs) were discovered, of which 243 were deemed specific interactors (spectral count greater than 4 in both WT animals, fold change greater than 2.0 between WT and KO). The STRING database^31,32^ matched 240 of these proteins which were subject to pathway and enrichment analysis using a medium confidence (0.4) threshold. Raw proteomics files and Scaffold file are available via MASSive (<u>doi:10.25345/C5CJ87Z98</u>).

### Brain tissue lysis and immunoblotting

Whole mouse striata were homogenized and sonicated in 1 mL in a low-ionic strength Tris buffer containing 2 mM Tris-HCl, 1 mM DTT, 2 mM EDTA, 1% Triton X-100, and 1X protease inhibitor cocktail (Bimake, Houston, TX, USA or APExBio, Houston, TX, USA), phosphatase inhibitors (20 mM sodium fluoride, 20 mM sodium orthovanadate, 20 mM β-glycerophosphate, and 10 mM sodium pyrophosphate; (Sigma-Aldrich or ThermoFisher Scientific Waltham, MA, USA)). Homogenates were incubated for 15 - 30 min at 4 °C and then centrifuged at 10,000 - 13,600 × *g* for 10 min. The cleared lysate was mixed with Laemmli sample buffer to generate input lysate samples. Striatal input lysates containing 1x Laemmli sample buffer with 250 µM 1,4-dithiothretol (DTT) (Sigma Aldrich, St. Louis, MO) were used for western blotting and all samples were initially heated at 70°C for 10 minutes. Each input sample (10-25 µg) was loaded into either a 1.5 mm hand-cast 10% polyacrylamide gel or a 26-well pre-cast Criterion 4-15% polyacrylamide gradient gel (Bio Rad Laboratories, Hercules, CA). The pre-cast gels were electrophoresed at 165 V for 1 hour, while the hand cast gels were electrophoresed at 75 V for 15 minutes and 165 V for 1 hour. Proteins were then transferred to a nitrocellulose membrane using either a wet transfer or the Trans-Blot Turbo semi-dry transfer (Bio-Rad Laboratories, Hercules, CA). For wet transfers, proteins were transferred to a nitrocellulose membrane using a N-cyclohexyl,1-3 aminopropanesulfonic acid (CAPS) transfer buffer (10% methanol, 0.01 M CAPS pH 11). The wet transfer was performed in a transfer tank attached to a cooling unit set at 4 °C and the transfer was performed at a constant 1.0 A for 1.5 hours. For the Trans-Blot Turbo transfer, gels were transferred to a nitrocellulose membrane using cold Trans-Blot Turbo buffer (Bio-Rad Laboratories, Hercules, CA) with 20% ethanol. The transfer was executed at a constant 1.0 amps for 30 minutes or 2.5 amps for 7 minutes. Following the transfer, membranes were stained with a 1 mg/mL Ponceau S stain dissolved in 10% Trichloroacetic acid or with a Revert 520 Total Protein Stain according to manufacturer’s instructions (LI-COR Biosciences, Lincoln, NE) to normalize for equal loading. Membranes were blocked with Tris-buffered saline with Tween (TBST: 50mM, Tris pH 7.5: 150 mM NaCl, 0.1% (volume/volume) Tween-20) containing 5% (weight/volume) nonfat dry milk. Blocking was performed 3 times, 10 minutes each, for a total of 30 minutes. Membranes were incubated with primary antibodies diluted 1:1000 in 5% milk in TBST overnight at 4 °C with gentle shaking. Primary antibodies used were mouse monoclonal anti-neurabin 1 (sc-136327, Santa Cruz Biotechnology Inc, Dallas, TX). After incubation, membranes were washed 3 times for 10 minutes per wash with TBST containing 5% milk. Appropriate secondary antibodies were typically diluted 1:10000 in TBST containing 5% milk. Appropriate secondary antibodies used were Alexa Fluor dye number 790 or 680 donkey anti-mouse (Intvitrogen, Waltham, MA). Membranes were incubated with the secondary antibody for 1 hour at room temperature, with gentle shaking in darkness. Membranes were washed in TBS without tween 3 times for 10 minutes each. Membranes were scanned using the Odyssey CLX or Odyssey M imaging system (LiCor, Lincoln, NE) and images were analyzed using Image Studio or Emperia Imaging Studio Software (LiCor, Lincoln, NE)

### Accelerating rotarod

A mouse rotarod accelerating from 4–40 rotations per minute over 300 seconds was used (**Figures 3, S2**: Rotamex-5, Columbus Instruments, Columbus, OH or **Figures 4**,**6**: 47650, Ugo Basile, Gemonio, Italy). Animals were trained with 3 trials per day for 5 days as previously described^26^. A trial ended when the mouse fell off the rotarod, they performed 2 consecutive tail rotations, or after 300 seconds had elapsed. Time to this endpoint was reported as “Latency to Fall”. Mice were allowed to rest 120-180 seconds between trials.

### Amphetamine Sensitization

Mice were intraperitoneally (IP) injected daily for 5 days with saline (Teknova, S5812), or 3 mg/kg d-amphetamine (Sigma-aldrich, A5880) dissolved in saline at a volume of 10 ml/kg. Immediately following each intraperitoneal (i.p.) injection, open field locomotion was recorded in Noldus Phenotyper Cages (Noldus Information Technology, Wageningen, Netherlands). Distance traveled and center time were recorded and quantified using Noldus XT software.

### Tamoxifen Treatment

Male and female Nrb^fl/fl^ and Nrb^fl/+^ animals expressing or lacking Cre recombinase under control of an estrogen receptor-dependent, CAG promoter (CAG-CreER) were treated with tamoxifen at 8 weeks of age. Animals were injected i.p. with 100 µL of 24 mg/mL tamoxifen solution every day for 5 days. Animals were sacrificed 3 weeks following the final injection and striatal tissue was collected and processed for analysis.

### RNAscope

Mice were transcardially perfused and brains were prepared for sectioning as previously described^26^. Ten micron striatal cryostat sections were mounted on Colorfrost Plus slides (#12-550-17, Thermo-Fisher Scientific) and taken through the RNAScope protocol for fixed frozen tissue samples as previously described^26^, using a target probe for neurabin (Mm-Ppp1r9a-C2, 1117781-C2, ACD) and the Opal 690 flourophore (#FP1497001KT, Akoya Biosciences, Marlborough, MA). To counterstain, slides were incubated with 4 drops of DAPI (#320858, ACD) for 30 seconds at RT and immediately incubated with ProLong Gold Antifade Mounting Mountant (#P36930, Thermo Scientific). Coverslips (#722204-01, Electron Microscopy Sciences, Hatfield, PA) were placed on tissue sections, sealed with nail polish, and allowed to dry overnight prior to imaging. Confocal fluorescence imaging for neurabin and DAPI was performed on the Zeiss LSM 900 with Airyscan 2. Dorsal striatal regions were chosen at the border of the corpus callosum and dorsal striatum and imaged using the PlanApochromat 63x/1.40 Oil objective, obtaining Z stacks (pixel size = 0.05 µm x 0.05 µm, step size = 0.13 micron). Z stacks were captured using Airyscan Multiplex 4Y functionality. Z-stacks (.czi files) were imported into ImageJ (National Institutes of Health; FIJI version 1.54p)^33^and a maximum intensity Z-projection of the individual channels was performed. To process neurabin channels, we performed background subtraction using FIJI’s *rolling ball* (50 px) plugin. Then, using levels derived from no probe batch controls, we set background/foreground image thresholding using the *Threshold* macro. Subsequently, neurabin puncta were quantified using Fiji’s *Analyze Particles* tool. Neurabin puncta were normalized to each ROI’s nuclei number obtained from a manual count in the DAPI channel. Representative images were generated by importing z projections into individual channels of an RGB image in Photoshop (Adobe) and adjusting levels linearly for picture presentation.

### Statistics

Statistical analysis was performed using GraphPad Prism 10.5.0 (GraphPad Software). Results are presented as the mean ± SEM. One outlier was excluded using a Grubb’s test. For each analysis, the statistical test used is reported in the figure legend. All statistics are reported in **Table S5**.

## Results

### Generation and validation of global neurabin knockout animals

PCR amplification around exon 6, the site of CRISPR targeting leading to non-homologous end joining, revealed the presence of a truncated gene product in neurabin heterozygote and knockout mice (**Figure 1A**). We confirmed a ~50% and complete knockout of neurabin in neurabin heterozygote and knockout mice, respectively (one-way ANOVA F(2,6)=156.1; p<0.0001, multiple comparisons +/+ vs. +/−; p=0.0001, multiple comparisons +/+ vs. −/−; p<0.0001, multiple comparisons +/−vs. −/−; p=0.0007) (**Figure 1B-C**). We characterized the baseline phenotype of our novel mouse line. Consistent with prior work suggesting a reduced anxiety phenotype observed in a previously established neurabin KO mouse line, neurabin global KO mice demonstrated increased center time in Noldus Phenotyper cages as compared to their WT counterparts (Unpaired T-test t=3.599 df=8, p=0.007) (**Figure 1D**)^23^. Compared to WT counterparts, male neurabin KO mice demonstrate decreased body mass at 4 weeks (Unpaired T-test, t=3.028, df=10; p=0.0127), but not 8 weeks (Unpaired T-test, t=1.382, df=10, p=0.1971) (**Figure 1E**), of age. By contrast, female neurabin KO mice display decreased body mass at both 4 weeks (Unpaired T test, t=2.581, df=8; p=0.0326) and 8 weeks (Unpaired T test, t=4.644, df=8; p=0.0017) (**Figure 1F**) of age. Finally, we assessed age-dependent expression of neurabin in the striatum. We found that neurabin is age dependently expressed, increasing steadily between PND 11 and 31, and peaking at 8 weeks of age (one-way ANOVA F(5,30)=8.946; p<0.0001, multiple comparisons P11 vs. P31; p=0.0014, P11 vs. 8 weeks; p<0.0001) (**Figure 1G-H**).

**Figure 1:**
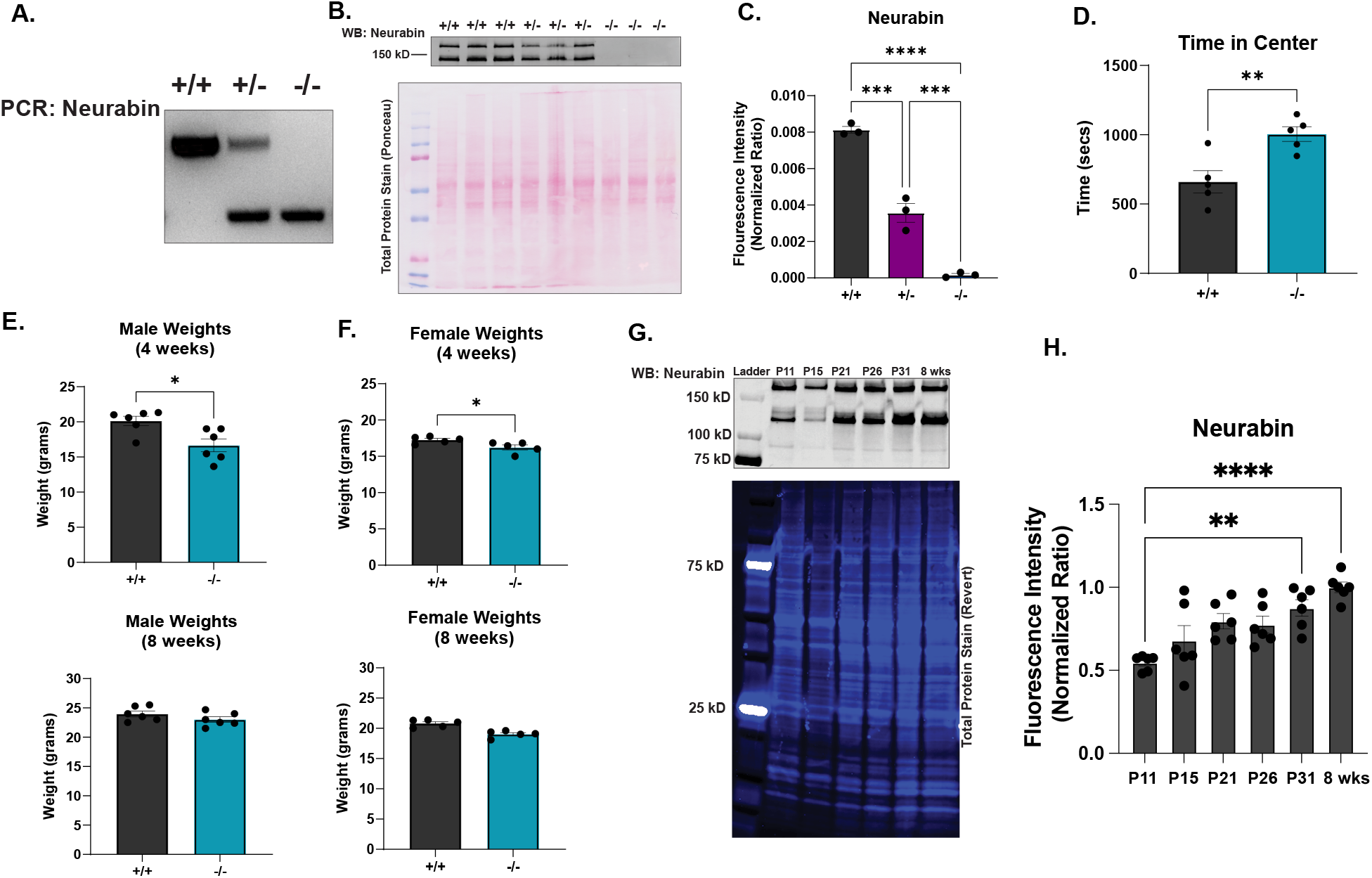
Generation and validation of global neurabin knockout animals. (**A**) DNA gel showing PCR products from neurabin wild type (Nrb^+/+^), heterozygote (Nrb^+/−^), and knockout (Nrb^−/−^) mice. (**B**) Neurabin immunoblot from Nrb^+/+^, Nrb^+/−^, and Nrb^−/−^ mice and Ponceau total protein stain. (**C**) One way ANOVA of Nrb^+/+^, Nrb^+/−^, and Nrb^−/−^ mice demonstrate a stepwise decrease in neurabin expression (one-way ANOVA F(2,6)=156.1; p<0.0001, multiple comparisons +/+ vs. +/−; p=0.0001, multiple comparisons +/+ vs. −/−; p<0.0001, multiple comparisons +/−vs. −/−; p=0.0007), n=3 per group, all male. (**D**) Time in center of saline treated animals on day 1 of sensitization experiment. Unpaired T-test demonstrates that Nrb^−/−^ mice spend more time in the center of area as compared to Nrb^+/+^counterparts (t=3.599 df=8, p=0.007), n=5 Nrb^+/+^ (2 male), n=5 Nrb^−/−^ (4 male) (**E**) Unpaired T-test demonstrates that 4 week old (t=3.028, df=10; p=0.0127) but not 8 week old (t=1.382, df=10, p=0.1971) male Nrb^−/−^ mice are lighter than their Nrb^+/+^ counterparts., n=6 per group (**F**) Unpaired T-test demonstrates that 4 week old (t=2.581, df=8; p=0.0326) and 8 week old (t=4.644, df=8; p=0.0017) week old female Nrb^−/−^ mice are lighter than their Nrb^+/+^ counterparts, n=5 per group (**G**) Neurabin immunoblot and Revert total protein stain of striata of Nrb^+/+^ mice dissected at P11, P15, P21, P26, P31, and 8 weeks of age. (**H**) One way ANOVA demonstrates that striatal neurabin expression is age dependent (F(5,30)=8.946; p<0.0001) and peaks at PND 31 and 8 weeks of age (multiple comparisons P11 vs. P31; p=0.0014, P11 vs. 8 weeks; p<0.0001)., n=6 P11 (3 male), n=6 P15 (4 male), n=6 P21 (3 male), n=6 P26 (3 male), n=6 P31 (4 male), n=6 8 week (4 male). Mean ± SEM. Significant ANOVA, T-Test, and post-hoc test results denoted by *p≤0.05, **p≤0.01, ***p≤0.001, ****p≤0.0001

### Connectivity and Function of Specific Neurabin Interacting Proteins

While neurabin interactions have been established from whole brain^34^, cortex^35^, and hippocampus^36–38^, or in vitro model systems^19,22,39^, the neurabin striatal interactome remains undefined. To comprehensively characterize the neurabin striatal interactome, we performed an unbiased analysis of the neurabin interactome using neurabin immunoprecipitation from WT or neurabin KO striatal lysates followed by mass spectrometry-based proteomics analysis. Out of the 1005 identified neurabin associated proteins (NrbAPs) (**Table S2A**), 243 were deemed specific (spectral count greater than 4 in both WT animals and a normalized fold change greater than 2.0 between WT and KO immunoprecipitates) (**Table S2B**). Then, 240 of these 243 proteins were matched in the STRING database and subject to pathway analysis (**Table S3A**, String-db). The top 10 “cellular component” GO pathways are enriched in proteins associated with the synapse, post synapse, glutamatergic synapse, and cell projection (**Figure 2A, Table S3B**). The top 10 “biological process” gene ontology (GO) pathways are enriched in proteins associated with organization of the actin cytoskeleton, synapse, post synapse, and cell junction (**Figure 2B, Table S3C**). The top 10 “molecular function” GO pathways are enriched in proteins associated with binding of cytoskeletal elements, protein complexes, enzymes, and anions (**Figure 2C, Table S3D**). The top 10 Kyoto Encyclopedia of Genes and Genome (KEGG) pathways are enriched in proteins associated with the tight junction, regulation of the actin cytoskeleton, neurological disorders, long term potentiation, and the glutamatergic synapse (**Figure 2D, Table S3E**). Because glutamatergic afferents comprise a major input to the striatum^3^ and affect GABAergic MSN output^40^, we elected to perform further analysis of the string interactome of those proteins matching the “glutamatergic synapse” cellular component GO term (**Figure 2E, Table S4**). Using String-db we identified 69 NrbAPs highlighted by “glutamatergic synapse” term and labeling this network with GO and KEGG terms (**Figure 2E**). We identified 13 NrbAPs uniquely associated with the postsynapse (*Adam 22, Anks1b, Rgs7*, etc.), modulation of chemical transmission (*Ppfia3*), and regulation of cellular component organization (*Amph, Rock2, Vcp*, etc.) (**Figure 2E**). The KEGG term “long term potentiation” (*Grin1-2a/b, Ppp1cc/a*, others) and GO term “structural components of the synapse” (*Actn1, Dlg1-4, Shank1-3*) were entirely composed of proteins associated with multiple GO and KEGG terms, characteristic of 53 of the remaining NrbAPs highlighted in this enrichment (**Figure 2E**).

**Figure 2:**
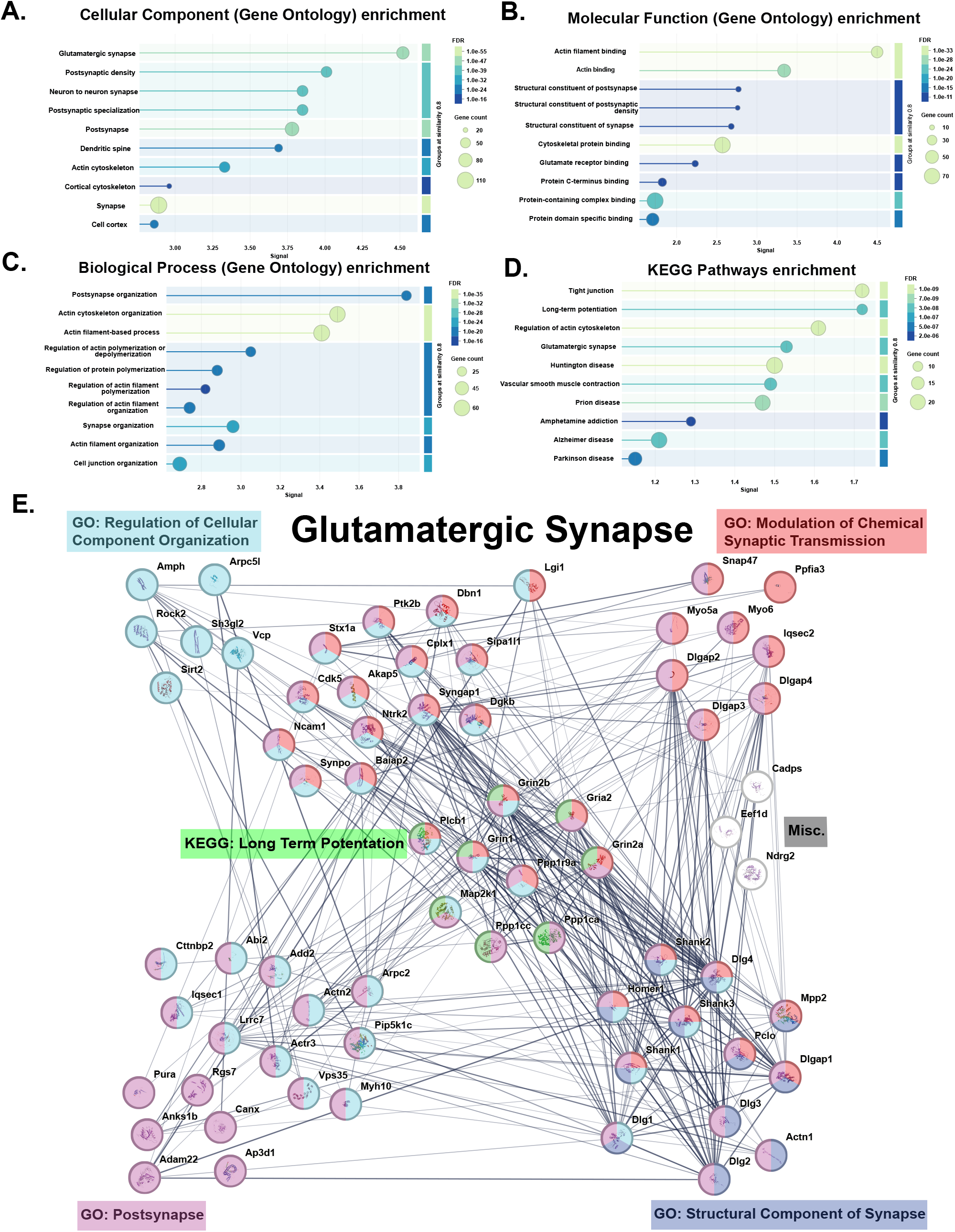
Connectivity and Function of Specific Neurabin Interacting Proteins. Specific neurabin interacting proteins based on peptide spectral matches (PSMs) were input into the string database (www.string-db.org). The database matched 240 total proteins. The false discovery rate (FDR), signal, and gene count for top 10 Gene Ontology (GO) enrichment terms based on signal matching **A**. “Cellular Component”, **B**. “Molecular Function, **C**. “Biological Process” as well as the **D**. Kyoto Encyclopedia of Genes and Genomes (KEGG) enrichment terms are shown. **E**. The proteins matching the “Glutamatergic Synapse” biological process and their string interactions are plotted. Within this enrichment, proteins matching the “Regulation of Cellular Component Organization”, “Modulation of Chemical Synaptic Transmission”, and “Long Term Potentiation”, “Postsynapse”, and “Structural Component of Synapse” were enriched. Proteins that do not match one of the pathways are labeled “Misc.”. Network edges indicate confidence interaction score, and thicker edges indicate greater stronger evidence of association. Medium confidence (0.4) used as lower cutoff.

### Young Adult Neurabin Global KO Mice Demonstrate Enhanced Rotarod Performance and Sensitize Normally to Amphetamine

Previous studies demonstrated that mice lacking spinophilin perform worse on an accelerating rotarod task ^20^ as compared to their WT counterparts and fail to sensitize to amphetamine^25^. However, the role of neurabin in motor learning and in psychomotor sensitization is not yet known. To assess these functions, a mixed sex cohort of 8-week-old littermate control (Nrb^+/+^) or whole body neurabin KO (Nrb^−/−^) were taken through a behavioral battery consisting of a 5-day accelerating rotarod paradigm, followed by a week of rest, and then 5 days of amphetamine sensitization (**Figure 3A**). The rotarod paradigm consisted of 3, 5-minute accelerating rotarod trials per day over 5 consecutive days. Latency to fall was recorded as a primary outcome measure and data was aggregated across days (**Figure 3B**) and individual trials (**Figure 3C**). While both groups exhibited motor learning, neurabin global KO mice broadly outperformed their littermate counterparts when analyzing data by day: Day (F(4,56)= 24.17; p<0.0001), Genotype (F(1, 14)=8.889; p=0.0099), Day x Genotype (F(4,56)=1.361; p=0.2592) or by individual trial: Trial (F(4.968,69.56)= 13.99; p<0.0001), Genotype (F(1, 14)=6.572; p=0.0225), Trial x Genotype (F(4.968,69.56)=0.8435; p=0.5230) (**Figure 3B-C**). Rotarod trained mice were subsequently administered either 3 mg/kg d-amphetamine or saline over 5 consecutive days and distance traveled was recorded as the primary outcome measure. While we observed treatment and day x treatment effects, demonstrating sensitization to amphetamine, there was no impact of genotype on locomotor responses to amphetamine. (Day (F(1.796, 21.56)=2.659; p=0.0975), Genotype (F(1,12)=0.4596; p=0.5106), Treatment (F(1,12=68.03); p<0.0001), Day x Genotype (F(1.796,21.56)=1.034; p=0.3653), Day x Treatment (F(1.796, 21.56) = 16.52; p<0.0001), Genotype x Treatment (F(1,12) = 0.09288; p=.7658), Day X Genotype X Treatment (F(1.796, 21.56)=0.9965; p=0.3776)) (**Figure 3D**).

**Figure 3:**
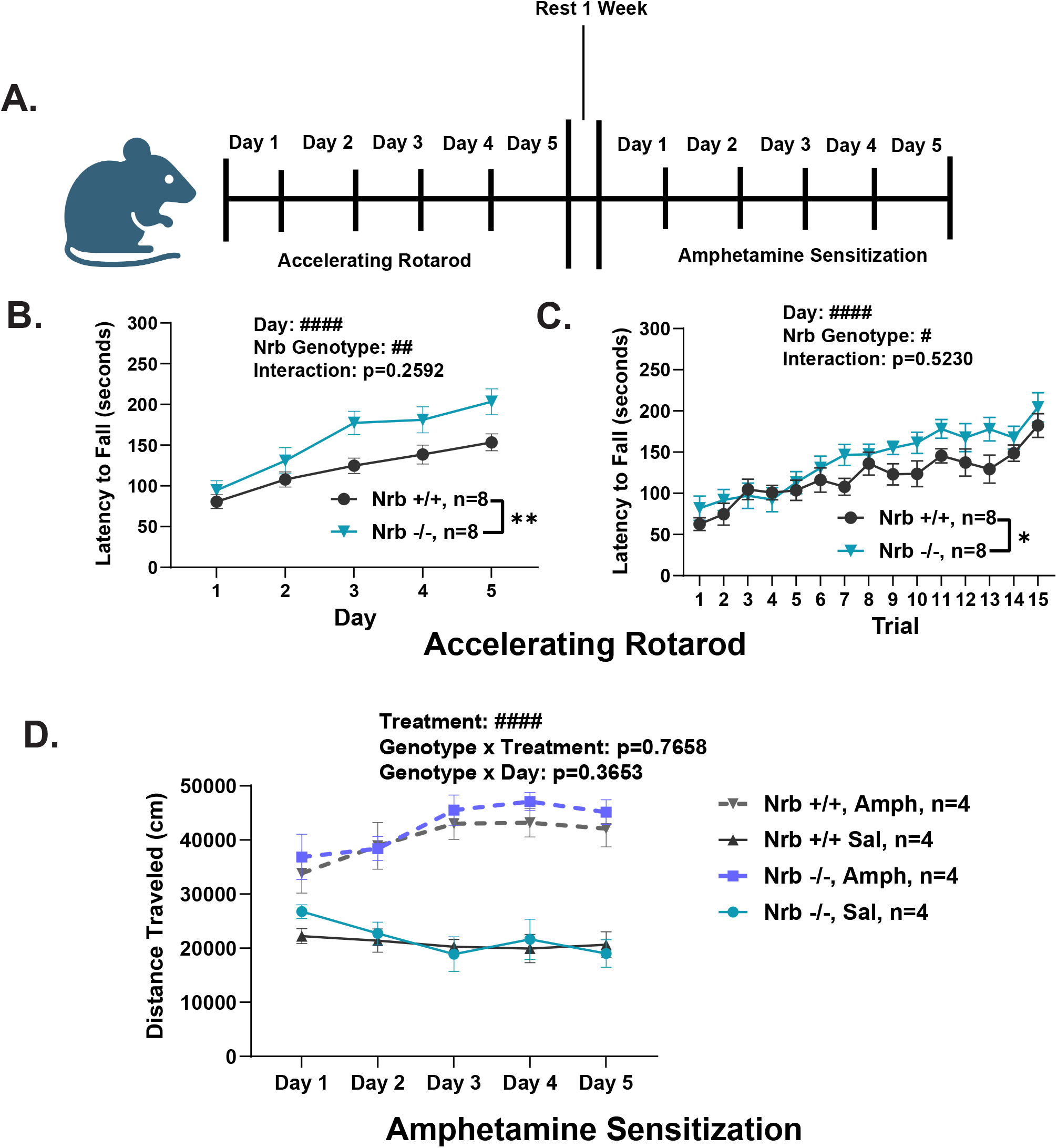
Young Adult Neurabin Global KO Mice Demonstrate Enhanced Rotarod Performance and Sensitize Normally to Amphetamine. (**A**) Mixed sex cohort of Neurabin WT (Nrb^+/+^) or Neurabin KO (Nrb^−/−^) mice taken through a behavioral battery. Mice are subject to a 5 consecutive day, 3 trial per day accelerating rotarod paradigm (4-40 RPM) followed by 1 week of rest, and 5 consecutive day amphetamine sensitization paradigm (3 mg/kg d-amphetamine or saline control) (**B**) Two way-RM ANOVA of Nrb^+/+^ and Nrb^−/−^ demonstrates that both cohorts learn the accelerating rotarod task (F(4,56)= 24.17; p<0.0001) but Nrb ^−/−^ outperform their littermate counterparts (F(1, 14)=8.889; p=0.0099) at the aggregate day level (**C**) Two way-RM ANOVA of Nrb^+/+^ and Nrb^−/−^ demonstrates that both cohorts learn the accelerating rotarod task (F(4.968,69.56)= 13.99; p<0.0001) but Nrb ^−/−^ outperform their littermate counterparts (F(1, 14)=8.889; p=0.0225) at the individual trial level. N=8 Nrb^+/+^ (3 male), n=8 Nrb^−/−^ (5 male) (**D**) Three-way ANOVAs with repeated measures detected significant Treatment (F(1,12)= 68.03; p<0.0001) and Day x Treatment (F(1.796,21.56)= 16.52; p<0.0001), but no Nrb^+/+^ or Nrb^−/−^ genotype or interaction effects on distance traveled daily injection with saline- or 3 mg/kg d-amphetamine. N=4/4 saline/AMPH-treated Nrb^+/+^ (1/2 male), 4/4 saline/AMPH-treated Nrb^−/−^ (3/2 male). Mean ± SEM. Significant ANOVA results denoted by #p≤0.05, ##p≤0.01, ###p≤0.001, ####p≤0.0001.

### Adult Male, but not Female, Neurabin Global KO Mice Demonstrate Enhanced Rotarod Performance

Convergent lines of evidence highlight sex specific differences in striatal physiology^41,42^, spine density^43^, and behavioral output^42^. Indeed, female rodents display enhanced locomotor activity^44^ and enhanced sensorimotor performance during estrus and in response to cannulated estradiol^45^. Accordingly, we sought to assess the sex specific impact of neurabin ablation in mature adult male and female mice. Eleven-to seventeen-week-old mice were taken through a 5-day accelerating rotarod paradigm as above. While male global KO and WT mice demonstrated motor learning, male global KO mice broadly outperformed their WT counterparts when analying data by day: Day (F(3.622,50.70)= 21.97; p<0.0001), Genotype (F(1, 14)=5.215; p=0.0385), and Day x Genotype (F(3.622,50.70)=2.128; p=0.0969)) or individual trial: Trial (F(6.215,87.01)= 10.70; p<0.0001), Genotype (F(1, 14)=5.215; p=0.0385), and Trial x Genotype (F(6.215,87.01)=1.457; p=0.2004) (**Figures 4A-B**). Female global KO and WT mice also demonstrated motor learning but did not perform differently from one another when analyzing data by day: Day (F(3.040,36.48)= 13.41; p<0.0001), Genotype (F(1, 12)=0.01244; p=0.9131), and Day x Genotype (F(3.040,36.48)=0.4290; p=0.7360) or by individual trial: Trial (F(5.454,65.45)= 9.571; p<0.0001), Genotype (F(1, 12)=0.009192; p=0.9252), and Trial x Genotype (F(5.454,65.45)=0.2645; p=0.9412) (**Figures 4C-D**). To assess whether neurabin was differentially expressed across age or sex in the mouse striatum in adult and mature animals, we immunoblotted striatal lysate from male and female mice at 8, 12, 16, and 44 weeks of age or older (**Figures 4E-F**). Adult neurabin expression was neither sex nor age dependent (Sex: F(1,39)=1.302; p=0.2609, Age: F(3, 39)=0.3327; p=0.8017, and Interaction: F(3, 39)=0.4304; p=0.7324).

**Figure 4:**
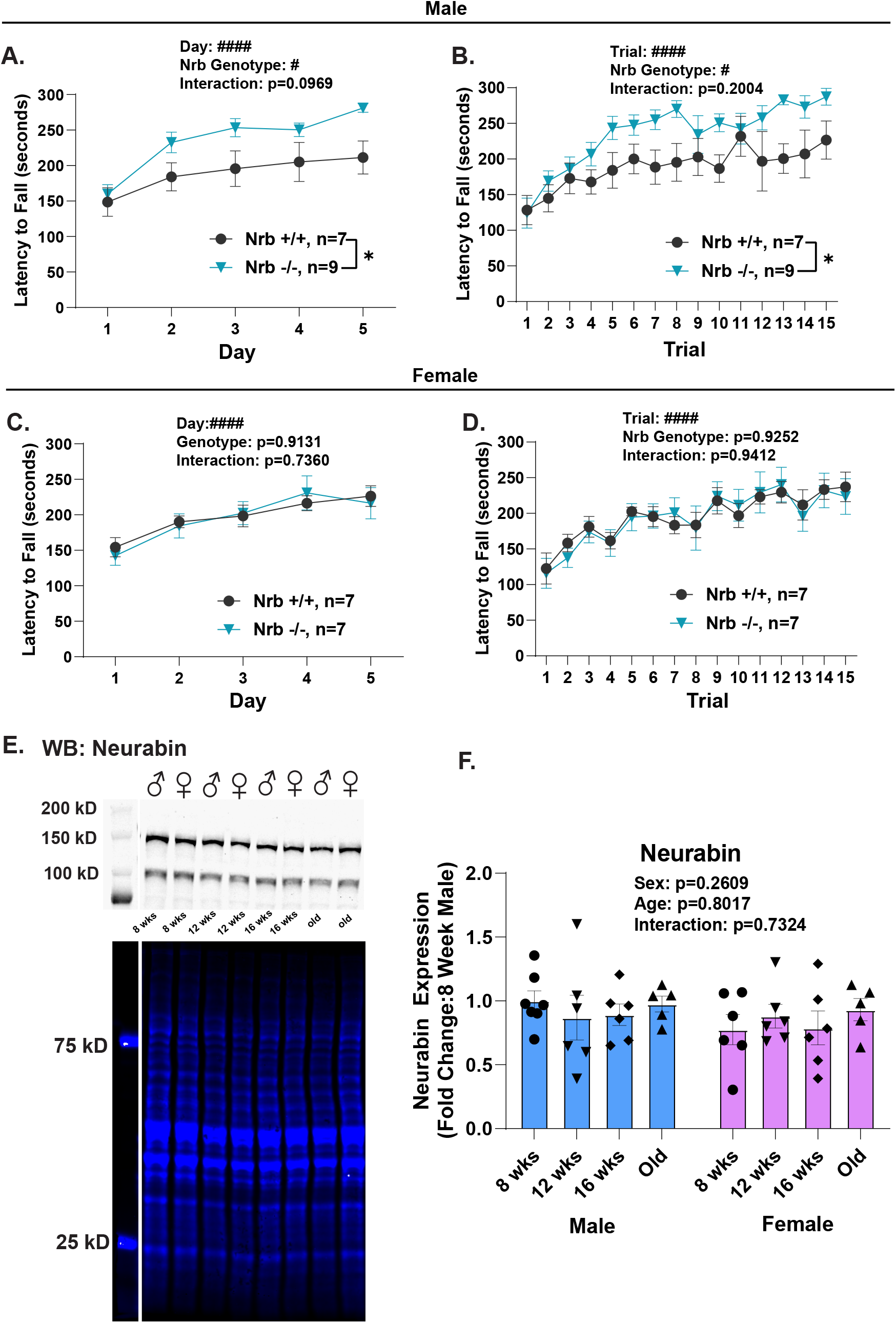
Young Adult Neurabin Global KO Mice Demonstrate Enhanced Rotarod Performance and Sensitize Normally to Amphetamine. (**A**) Two way-RM ANOVA of male Nrb^+/+^ and Nrb^−/−^ mice demonstrates that both cohorts learn the accelerating rotarod task (F(3.622,50.70)= 21.97; p<0.0001) but Nrb ^−/−^ mice outperform their littermate counterparts (F(1, 14)=5.215; p=0.0385) at the aggregate day level (**B**) Two way-RM ANOVA of male Nrb^+/+^ and Nrb^−/−^ demonstrates that both cohorts learn the accelerating rotarod task (F(6.215,87.01)= 10.70; p<0.0001) but Nrb ^−/−^ outperform their littermate counterparts (F(1, 14)=5.215; p=0.0385) at the individual trial level. (**C**) Two way-RM ANOVA of female Nrb^+/+^ and Nrb^−/−^ demonstrates that both cohorts learn the accelerating rotarod task (F(3.040,36.48)= 13.41; p<0.0001) but Nrb ^+/+^ and Nrb^−/−^ perform similarly to their littermate counterparts (F(1, 12)=0.01244; p=0.9131) at the aggregate day level. (**D**) Two way-RM ANOVA of female Nrb^+/+^ and Nrb^−/−^ demonstrates that both cohorts learn the accelerating rotarod task (F(5.454,65.45)= 9.571; p<0.0001) but Nrb ^+/+^ and Nrb^−/−^ perform similarly to their littermate counterparts (F(1, 12)=0.009192; p=0.9252) at the individual trial level. (**E**) Neurabin immunoblot and Revert total protein stain of striata of male and female Nrb^+/+^ mice dissected at 8 weeks, 12 weeks, 16 weeks, and 44+ weeks of age (old). Two way ANOVA demonstrates that striatal neurabin expression does not vary by age (F(3, 39)=0.3327,p=0.8017), sex (F(1,39)=1.302; p=0.2609), or the interaction of age or sex (F(3, 39)=0.4304; p=0.7324). n=5-7 per sex per group. Mean ± SEM. Significant ANOVA results denoted by #p≤0.05, ##p≤0.01, ###p≤0.001, ####p≤0.0001.

### Generation and Validation of Cell Type-Specific Neurabin KO Mice

Previous studies have demonstrated that ablation of spinophilin impairs rotarod motor learning in an iMSN and dMSN specific manner^26^, and that dMSNs and iMSNs individually and synergistically contribute to motor coordination and motor learning^46,47^. To examine the cell type specific effects of neurabin, we developed a conditional neurabin allele by inserting loxp sites around exon 6. This was done in collaboration with the Transgenic Animal Model Core of the University of Michigan’s Biomedical Research Core Facilities. To validate the flox dependent Cre recombinase-mediated depletion of neurabin, we crossed our neurabin flox mice with CAG-CreER mice to generate a tamoxifen inducible, Cre recombinase-mediated conditional neurabin knockout mouse (Nrb^CagCreER^). Following tamoxifen injection, Nrb^CagCreER^ mice demonstrated stepwise neurabin depletion in Nrb^fl/fl^/Cre-, Nrb^fl/^+/Cre+, and Nrb^fl/fl^/Cre+ genotypes (One-Way ANOVA, F=7.943; p=0.0158, multiple comparisons fl/fl Cre-vs. fl/fl Cre+; p=0.0105, fl/fl Cre-vs. fl/+ Cre+; p=0.0733) (**Figure 5A-B**). To generate cell type specific neurabin knockout mice, we crossed Nrb^fl/fl^ mice with mice expressing cre recombinase under control of either the Drd1 (Cre^D1^) or Adora2a (Cre^A2a^) genes to express cre in dMSNs (Nrb^ΔdMSN^) or iMSNs (Nrb^ΔiMSN^) similar to the conditional spinophilin line previously generated by our lab^26^. To assess striatal neurabin depletion, we performed RNAScope on striatal sections. When examining ROIs of the dorsal striatum, we observed a qualitative, but not statistical decrease in the number of neurabin mRNA puncta in Nrb^ΔdMSN^ (Unpaired T test, t=2.213, df=4; p=0.0914) and Nrb^ΔiMSN^ (Unpaired T test, t=1.723, df=4; p=0.1601) mice relative to their Nrb^fl/fl^ counterparts (**Figure 5C**). This amounted to a ~40% depletion of neurabin mRNA that was statistically different in Nrb^ΔdMSN^ (Unpaired T test, t=3.715, df=4; p=0.0206) but not Nrb^ΔiMSN^ mice relative to their Nrb^fl/fl^ counterparts (Unpaired T test, t=1.728, df=5; p=0.159) (**Figure 5D**). Furthermore, we observed a selective depletion of neurabin from some but not all striatal nuclei in the Nrb^ΔdMSN^ and Nrb^ΔiMSN^ mice relative to Nrb^fl/fl^ counterparts (**Figure 5E, 5F**).

**Figure 5:**
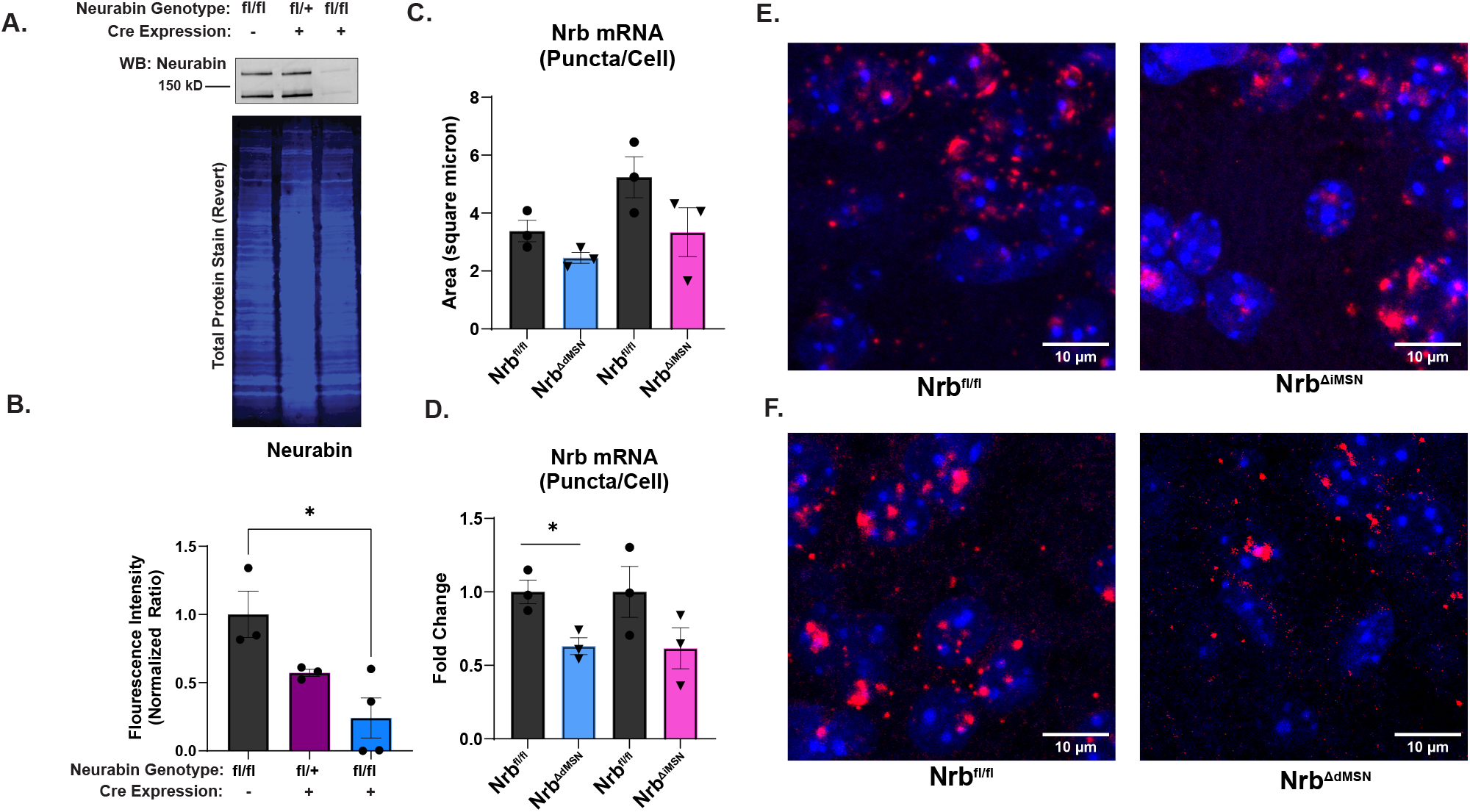
Generation and Validation of Neurabin Conditional and Cell Type Specific Neurabin Knockout Mice. (**A**) Neurabin immunoblot and Revert total protein stain of striata of male and female mice of Nrb^CagCreER^, Nrb^fl/fl^, and Nrb^fl/+^/CagCre^+^ subjected to 5 consecutive days of Tamoxifen treatment at 8 weeks of age. Striata harvested at 11 weeks. (**B**) One way ANOVA demonstrates CagCre significantly depletes striatal neurabin (F=7.943; p=0.0158, multiple comparisons fl/fl Cre-vs. fl/fl Cre+; p=0.0105), and trends towards a stepwise depletion across fl/fl and fl/+ genotypes (multiple comparisons fl/+ Cre+ vs. fl/fl Cre+; p=0.0733) n=4 (3 male) Nrb^CagCreER^ (3 male), n=3 Nrb^fl/fl^ (2 male), and n=3 Nrb^Fl/+^/CagCre^+^ (3 female) (**C**) Nrb^fl/fl^ mice were crossed with Nrb^D1Cre^ and Nrb^A2aCre^ mice to make mice cell type specifically lacking neurabin in dMSNs (Nrb^ΔdMSN^) and iMSNs (Nrb^ΔiMSN^). To validate neurabin mRNA depletion within the striatum, we performed RNAscope of neurabin in coronal sections originating from male and female Nrb^ΔdMSN^ or Nrb^ΔiMSN^ mice and Nrb^fl/fl^ mice. Three ROIs spanning the dorsal striatum were imaged per mouse and neurabin puncta were normalized to nuclei number within each ROI. Within animals of the same condition, Nrb mRNA values from anatomically matching ROIs were grouped and compared in aggregate between Nrb^ΔdMSN^ or Nrb^ΔiMSN^ mice and their flox littermate or age matched counterparts. Unpaired T-tests demonstrate that compared to their flox counterparts, Nrb^ΔdMSN^ (t=2.213, df = 4, p=0.0914) and Nrb^ΔiMSN^ (t=1.723, df=4, p=0.1601) mice trend towards lower mean nrb puncta/cell. (**D**) Fold change values were obtained by comparing average intensity per ROI per mouse to the average mRNA intensity across ROIs in littermate or age matched flox controls. Unpaired T-tests demonstrate that Nrb^ΔdMSN^ (t=3.715, df = 4, p=0.0206) and Nrb^ΔiMSN^ (t=1.728, df=4, p=0.159) mice show or trend towards lower nrb mRNA/cell, amounting to ~40% reduction in nrb mRNA intensity relative to matched flox controls. N=2 Nrb^ΔdMSN^ (1 male), n=2 Nrb^ΔiMSN^ (1 male), n=4 Nrb^fl/fl^ (2 male) (**E**) Nrb^ΔdMSN^ and Nrb^ΔiMSN^ mice demonstrate cell specific depletion of neurabin as compared to their Nrb^fl/fl^ counterparts. Mean ± SEM. Significant ANOVA, T-Test, and post-hoc test results denoted by *p≤0.05, **p≤0.01, ***p≤0.001, ****p≤0.0001

### Ablation of Neurabin Sex and Cell Type Specifically Enhances Accelerating Rotarod Performance

To assess neurabin’s cell type specific functions in motor learning,10-15-week-old control (Nrb^fl/+^, Nrb^fl/fl^, Nrb^+/+^/Cre^D1^, or Nrb^+/+^/Cre^A2a^), Nrb^ΔdMSN^, and Nrb^ΔiMSN^ mice were taken through an accelerating rotarod paradigm as prior. While male control and Nrb^ΔdMSN^ mice demonstrated motor learning, male Nrb^ΔdMSN^ mice outperformed control mice when analyzed by individual day: Day (F(4, 92) = 32.86; p<0.0001), Genotype (F(1, 23) = 6.344; p=0.0192), Day x Genotype (F(4, 92)=0.1755; p=0. 9505)) and or by individual trial: Trial (F(7.482, 172) = 15.64; p<0.0001), Genotype (F(1, 23) = 6.293; p=0.0196), Trial x Genotype (F(7.482, 172)=0.3731; p=0.9256) (**Figure 6A-B**). While female control and Nrb^ΔdMSN^ mice demonstrated motor learning, female Nrb^ΔdMSN^ mice performed similarly to control mice when analzyed by individual day: Day (F(3.235, 58.24) = 17.98; p<0.0001), Genotype (F(1, 18) = 0.4500; p=0.5180), Day x Genotype (F(3.235, 58.24)=0.6849; p=0.5755) or by individual trial: Trial (F(7.376, 132.8) = 13.97; p<0.0001), Genotype (F(1, 18) = 0.3161; p=0.5809), Trial x Genotype (F(7.376, 132.8)=0.1.025; p=0.4182) (**Figure 6C-D**).

**Figure 6:**
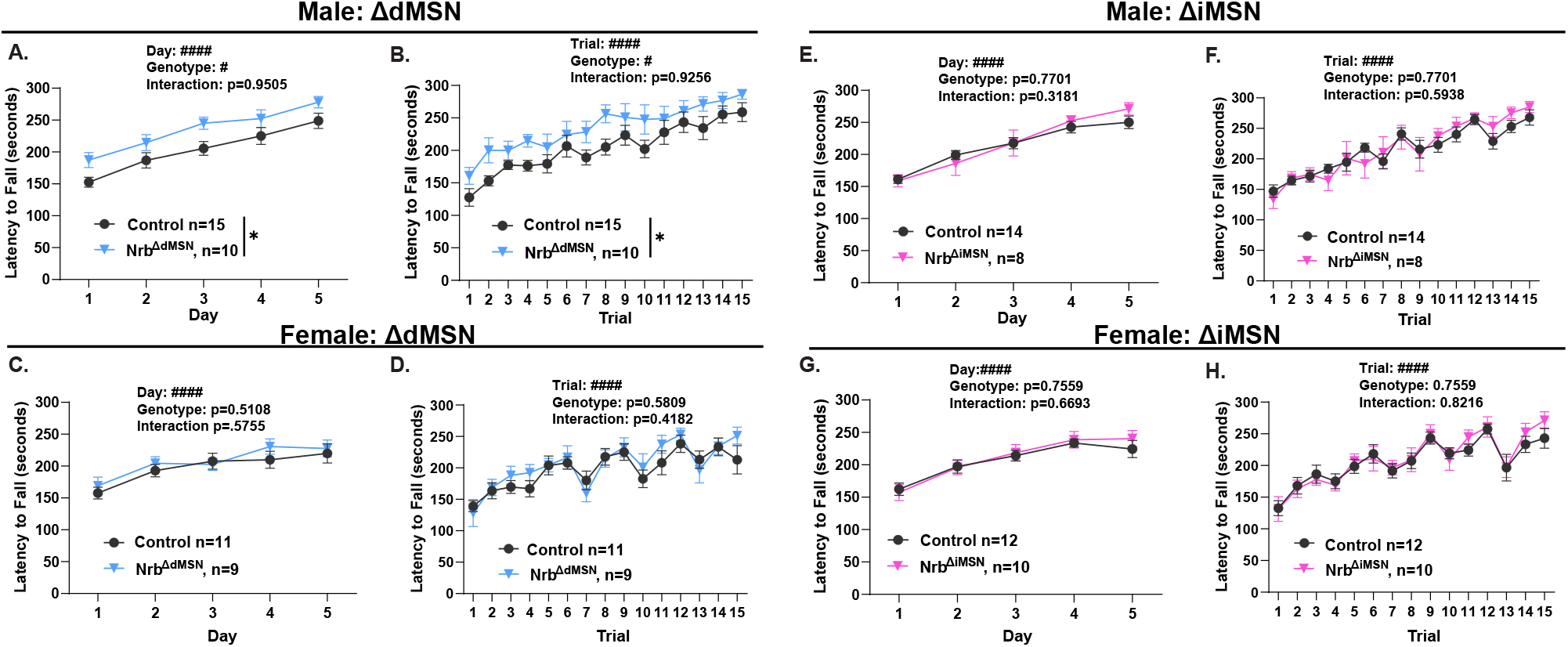
Ablation of Neurabin Sex and Cell Type Specifically Enhances Accelerating Rotarod Performance. (**A**) Two way-RM ANOVA of male Nrb^ΔdMSN^ and Nrb^fl/fl^ or Nrb^+/+^/Cre^D1^ mice demonstrates that both cohorts learn the accelerating rotarod task (F(4, 92) = 32.86; p<0.0001) but Nrb^ΔdMSN^ mice outperform Nrb^fl/fl^ or Nrb^+/+^/Cre^D1^ counterparts (F(1, 23) = 6.344; p=0.0192) at the aggregate day level (**B**) Two way-RM ANOVA of male Nrb^ΔdMSN^ and Nrb^fl/fl^ or Nrb^+/+^/Cre^D1^ mice demonstrates that both cohorts learn the accelerating rotarod task (F(7.482, 172) = 15.64; p<0.0001) but Nrb^ΔdMSN^ mice outperform Nrb^fl/fl^ or Nrb^+/+^/Cre^D1^ counterparts (F(1, 23) = 6.344; p=0.0196) at the aggregate trial level (**C**) Two way-RM ANOVA of female Nrb^ΔdMSN^ and Nrb^fl/fl^ or Nrb^+/+^/Cre^D1^ demonstrates that both cohorts learn the accelerating rotarod task (F(3.235, 58.24) = 17.98; p<0.0001) but female Nrb^ΔdMSN^ and Nrb^fl/fl^ or Nrb^+/+^/Cre^D1^ perform similarly (F(1, 18) = 0.4500; p=0.5108) at the aggregate day level. (**D**) Two way-RM ANOVA of female Nrb^ΔdMSN^ and Nrb^fl/fl^ or Nrb^+/+^/Cre^D1^ demonstrates that both cohorts learn the accelerating rotarod task (F(7.376, 132.8) = 13.97; p<0.0001) but female Nrb^ΔdMSN^ and Nrb^fl/fl^ or Nrb^+/+^/Cre^D1^ perform similarly (F(1, 18) = 0.3161; p=0.5809) at the individual trial level. (**E**) Two way-RM ANOVA of male Nrb^ΔiMSN^ and Nrb^fl/fl^ or Nrb^+/+^/Cre^A2a^ demonstrates that both cohorts learn the accelerating rotarod task (F(2.846, 56.93) = 48.07; p<0.0001) but male Nrb^ΔiMSN^ and Nrb^fl/fl^ or Nrb^+/+^/Cre^A2a^ perform similarly (F(1, 20)= 0.08779; p=0.7701) at the aggregate day level. (**F**) Two way-RM ANOVA of male Nrb^ΔiMSN^ and Nrb^fl/fl^ or Nrb^+/+^/Cre^A2a^ demonstrates that both cohorts learn the accelerating rotarod task (F(6.572, 131.4) = 22.53; p<0.0001) but male Nrb^ΔiMSN^ and Nrb^fl/fl^ or Nrb^+/+^/Cre^A2a^ perform similarly (F(1, 20) =0.08779; p=0.7701) at the individual trial level. Two way-RM ANOVA of female Nrb^ΔiMSN^ and Nrb^fl/fl^ or Nrb^+/+^/Cre^A2a^ demonstrates that both cohorts learn the accelerating rotarod task (F(2.891, 57.82) = 30.88; p<0.0001) but female Nrb^ΔiMSN^ and Nrb^fl/fl^ or Nrb^+/+^/Cre^A2a^ perform similarly (F(1, 20) =0.09935; p=0.7559) at the aggregate day level. (**H**) Two way-RM ANOVA of female Nrb^ΔiMSN^ and Nrb^fl/fl^ or Nrb^+/+^/Cre^A2a^ demonstrates that both cohorts learn the accelerating rotarod task (F(6.509, 138.1) = 20.77; p<0.0001) but female Nrb^ΔiMSN^ and Nrb^fl/fl^ or Nrb^+/+^/Cre^A2a^ perform similarly (F(1, 20) = 0.009935; p=0.7559) at the individual trial level. Mean ± SEM. Significant ANOVA results denoted by #p≤0.05, ##p≤0.01, ###p≤0.001, ####p≤0.0001.

We also evaluated loss of neurabin in iMSNs. In contrast to loss in dMSNs, male control and Nrb^ΔiMSN^ mice demonstrated motor learning but performed similarly to each other when analyzing by day: Day (F(2.846, 56.93) = 48.07; p<0.0001), Genotype (F(1, 20) = 0.08779; p=0.7701), Day x Genotype F(2.846, 56.93)=1.197; p=0.3181) or by individual trial: Trial (F(6.572, 131.4) = 22.53; p<0.0001), Genotype (F(1, 20) = .08779; p=0.7701), Trial x Genotype (F(6.572, 131.4)=0.7849; p=0.5938) (**Figure 6E-F**). Female control and Nrb^ΔiMSN^ mice also demonstrated similar motor learning to each other when analyzing by day: Day (F(2.891, 57.82) = 30.88; p<0.0001), Genotype (F(1, 20) = 0.09935; p=0.7559), Day x Genotype (F(2.891, 57.82)=0.5114; p=0.6693) or by trial: Trial (F(6.509, 138.1) = 20.77; p<0.0001), Genotype (F(1, 20) = 0.009935; p=0.7559), Trial x Genotype (F(6.905, 138.1)=0.5126; p=0.8216) (**Figure 6G-H**).

## Discussion

Neurabin and its homolog spinophilin were initially characterized as actin binding proteins and the two most-abundant PSD-enriched PP1 binding proteins^34,48^. Despite their homology and overlapping functions, prior studies have delineated distinct phenotypic and biochemical roles for both proteins, particularly in their roles in corticostriatal plasticity^24^. Spinophilin has been demonstrated to enforce corticostriatal LTD^24^, which occurs in a dopamine D2 receptor-dependent manner^49^. By contrast, neurabin has been demonstrated to enforce corticostriatal LTP^24^ which occurs in an NMDAR-dependent manner^50^. While the spinophilin striatal interactome has been well characterized^51–53^, the neurabin striatal interactome is less well defined.

To characterize the neurabin interactome, we performed an unbiased proteomics screen of neurabin immunoprecipitates isolated from striatal lysates. Our proteomic analysis of the striatal neurabin interactome highlighted “glutamatergic synapse” as one of the most enriched (by false discovery rate) gene ontology (GO) terms within the Cellular Component pathway. Within the “glutamatergic synapse” cellular component term, we observed NMDAR subunits (*Grin1, Grin2a, Grin2b* genes) which play critical roles in LTP induction^54,55^. We also identified proteins like *Map2k1* and *Gria2* genes which serve to stabilize NMDAR LTP^56,57^, alongside *Plcb1, Ppp1ca, Ppp1cc*, which function in homeostatic roles within signaling cascades to balance NMDAR LTP in the striatum^58,59^. We also identified several scaffolding proteins of the MAGUK (*Mpp2, Dlg1-4*), SAPAP (*Dlgap1-4*) and SHANK (*Shank1-3*), and Homer (*Homer1*) families which serve critical roles in the stabilization of synaptic AMPA and NMDA receptors^60^, in synaptogenesis and synaptic scaling^61,62^, and in bridging ionotropic and metabotropic activity at glutamate synapses^63^, respectively. These results are consistent with prior findings that ablation of neurabin abolishes corticostriatal LTP^24^ and that neurabin facilitates the outgrowth of filopodia^64^. They are also consistent with the finding that neurabin/PP1 interactions limit the outgrowth of filopodia in cultured neurons^65^ and the neurabin/PP1 interaction regulates synaptic GluA1 and GluA2 trafficking^64,66^. In the striatum, neurabin ablation decreases levels of PP1 and increases phosphorylation of S845 on GluA1 in response to the dopamine D1 agonist, SKF 81297^24^, suggesting that neurabin limits dopamine mediated insertion of AMPA receptors into the PSD^57^.

As expected, we identified several actin cytoskeleton associated proteins like *Aprc5l*^64,67^ and endophilin which modulates spine morphology and plasticity-dependent actin polymerization^65^. While Arp2/3 and endophilin can impact post synaptic physiology^64,66^, endophilin also impacts presynaptic physiology, alongside amphiphysin and piccolo, by regulating synaptic vesicle trafficking ^68^. While most of the neurabin interactome was post-synaptic (**Figure S1**), the identification of proteins with pre- and post-synaptic roles is consistent with subcellular localization studies that find neurabin is also present in dendrites, axons and axon terminal^17^. Furthermore, the presence of presynaptic vesicle trafficking proteins in our proteomics is congruent with the finding that neurabin ablation increases the frequency of miniature and spontaneous excitatory postsynaptic currents in striatal neurons^24^ and suggests that neurabin impacts elements of presynaptic physiology and limits neurotransmitter release. Collectively, these data present a putative role for neurabin in the induction and modulation of LTP and the development, maintenance, and modulation of glutamatergic synapse function.

Glutamatergic synapses arise from thalamic and cortical projections into the striatum^1^, and plasticity at these synapses underlies striatal function^3^. Accordingly, we implemented a behavioral battery to examine the role of neurabin in 2 striatal behaviors associated with changes in striatal plasticity: rotarod motor learning^46^ and amphetamine sensitization^69^. Global ablation of neurabin increased rotarod performance in a mixed cohort of 8-week-old male and female mice as compared to WT controls. Interestingly, amphetamine sensitization was similar between global neurabin knockout mice and WT controls, suggesting that while neurabin impacts glutamate driven skill motor learning^46^, it doesn’t impact motor sensitization associated with psychostimulant-induced behavioral plasticity^69^. While we were not powered to detect within or between sex effects in this cohort, our data suggested that the effect of neurabin ablation may be more robust in female mice at this age (**Figure S2**).

Given that sex differences impact striatal physiology and behavioral output^41–45^, we examined the impact of neurabin ablation in adult male and female mice. Interestingly, male, but not female, global neurabin knockout mice outperformed their WT counterparts on the accelerating rotarod task in adult mice. Furthermore, we found that from 8 weeks onward, striatal expression of neurabin didn’t differ across age or sex, suggesting that neurabin-dependent limitation on skill motor learning may be subject to overarching sex specific regulation. Because striatal dMSNs and iMSNs play distinct but synergistic roles in rotarod motor learning^46,70,71^, we generated and validated cell type-specific neurabin KO animals lacking neurabin in dMSNs or iMSNs. We then trained cohorts of Nrb^ΔdMSN^ and Nrb^ΔiMSN^ adult mice on the accelerating rotarod task as before. Nrb^ΔdMSN^, but not Nrb^ΔiMSN^, mice outperformed their WT counterparts on a rotarod motor learning task. While impairment or ablation of iMSNs has been demonstrated to dramatically limit performance and learning on the rotarod motor learning task^46,70,71^, dMSNs uniquely underlie the performance improvements in the accelerating rotarod motor learning paradigm^47^, promote the early stage development of motor coordination^72^ and promote the initiation and perseverance in a motor learning task generally^2^. Congruently, our Nrb^ΔdMSN^ demonstrate immediate and persistent performance enhancement on the accelerating rotarod task as compared to their floxed counterparts, suggesting that dMSN specific ablation of neurabin may enhance direct pathway output. As we observed in our global neurabin KO animals, this effect was observed in male, but not female, mice furthering the idea that the effect of neurabin is subject to sex specific regulation even within striatal cell types.

As expected for homologous proteins, spinophilin and neurabin share sequence similarity and PP1 targeting roles^15,16^, overlapping subcellular localizations^17^, both regulate dendritic spine morphology^21,22,73^, and co-immunoprecipitate in the striatum^51^. Prior work has delineated the spinophilin striatal interactome^26,51,53,74^, demonstrated that spinophilin ablation impairs rotarod motor learning^20^ through its role in iMSNs^26^, and impairs amphetamine sensitization^25^. Our findings demonstrate that while the neurabin striatal interactome largely overlaps with that of spinophilin^26,51,53,74^, neurabin ablation enhances rotarod motor learning through its actions in dMSNs and does not impact amphetamine sensitization. The overlap in the interactomes of these two proteins suggests that spinophilin’s GPCR domain^75^, and spinophilin’s interactions with mglur5^76^ and the dopamine D2 receptor^77^ may underlie the functional difference in both proteins. Furthermore, while spinophilin is locally translated at the synapse, neurabin mRNA is found in the soma, consistent with cell body translation^78,79^. These findings, taken alongside the finding that neurabin protein is found throughout the cell and is both pre and post-synaptic, contrasted with spinophilin, which is enriched in the post-synapse^17^, suggest that while the interactomes may be similar, the actions of both proteins may be modified by the spatial context of the different neuronal compartments they are synthesized and reside in.

## Conclusions and Future Studies

Our data present a comprehensive striatal neurabin interactome and leverage novel neurabin mouse models to delineate the global impact of neurabin on amphetamine sensitization and the global and cell type specific impact of neurabin on accelerating rotarod motor learning. Broadly, a notable subset of neurabin interacting proteins are known to function within the glutamatergic synapse, playing roles in the induction and modulation of LTP, as well as the development and maintenance of glutamatergic synapse structure and function. Furthermore, ablation of neurabin enhances performance on the accelerating rotarod skill motor learning task but does not impact amphetamine psychomotor sensitization. While this effect was initially discovered in adolescent (8-week-old) mixed sex cohort of male and female mice, when examined in adult animals, neurabin ablation was found to enhance rotarod performance in males but not females. This finding was recapitulated in our male neurabin dMSN knockout mice, demonstrating that neurabin sex- and cell type-specifically limits skill motor learning in adult mice. While future studies need to delineate how skill motor learning impacts cell type-specific neurabin protein interactions and how loss of neurabin impacts cell type-specific synaptic protein phosphorylation and expression changes caused by motor learning, this current study is the first to present a comprehensive neurabin striatal interactome and characterize neurabin as a regulator of striatal-associated motor performance. These findings expand a growing literature that delineates unique biochemical, proteomic, and behavioral roles for the homologous actin binding proteins, spinophilin and neurabin. They also suggest novel cell type specific mechanisms by which scaffolding proteins and PSD biology can regulate striatal physiology and striatal-dependent motor function.

## Acknowledgments

We acknowledge and thank Wanda Filipiak & Galina Gavrilina for embryo injections for the initial generation of Neurabin^fl/fl^ mice as well as the entire excellent Transgenic Animal Model Core past and present (in particular, Anna LaForest, Elizabeth Hughes, Corey Ziebell, Dr. Thomas Saunders, and Dr. Zach Freeman) and the University of Michigan’s Biomedical Research Core Facilities for their generation of these mice. We acknowledge Drs. Emma Doud and Amber Mosley in the Indiana University School of Medicine Center for Proteome Analysis for technical support and expertise for mass spectrometry. Acquisition of the IUSM Center for Proteome Analysis instrumentation used for this project was provided by the Indiana University Precision Health Initiative. The proteomics work was supported, in part, by the Indiana Clinical and Translational Sciences Institute (UL1TR002529 from the National Institutes of Health, National Center for Advancing Translational Sciences, Clinical and Translational Sciences Award) and the P30 IU Simon Comprehensive Cancer Center Support Grant (P30CA082709). Funding for the generation of these mice and completion of studies comes from an R21/R33 award from the National Institutes of Drug Abuse (R21/R33 DA041876 to AJB), Department of Biology/School of Science at IU-Indy, Department of Pharmacology and Toxicology Startup Funds, and Strategic Research Initiative Funds (Indiana University School of Medicine and Stark Neurosciences Research Institute).

