## Supplementary material for "Proteomic Characterization of Striatal Neurabin Interactome and its Sex Specific Impact on Motor Behavior": Figure S

**Supplemental Information.**

**Table S1. Primer Sequences for Genotyping Experimental Mouse Lines.**

**Table S2: Neurabin Interactome.** **A.** List of all proteins detected by mass spectrometry and their spectral counts in neurabin immunoprecipitates isolated from 2 WT and 2 KO tissue samples. **B.** List of specific neurabin interacting proteins based on normalized (to total spectra) fold-change > 2 between one-pair of WT and KO samples and 4 or more peptide spectral matches. Known contaminants were removed.

**Table S3. Protein Descriptors and Pathway Analysis for Neurabin Striatal Interactome. A.** Alphabetical gene name and protein alias and descriptions for the 240 proteins that were specific neurabin interactors and present in the STRING-db. **B.** Enrichment for the “Cellular Component” GO term arranged by false discovery rate (FDR). Includes list of matching proteins in each network. **C.** Enrichment for the “Molecular Function” GO term arranged by false discovery rate (FDR). Includes list of matching proteins in each network. **D.** Enrichment for the “Biological Process” GO term arranged by false discovery rate (FDR). Includes list of matching proteins in each network. **E.** Enrichment for the Kyoto Genes and Genomes pathway arranged by false discovery rate (FDR). Includes list of matching proteins in each network.

**Table S4: Protein Descriptors and Pathway Analysis for the "Glutamatergic Synapse" Enrichment within the “Cellular Component” GO Term from the Specific Neurabin Striatal Interactome. A.** Alphabetical gene name and protein alias and descriptions for the 69 proteins that were specific neurabin interactors matching the Glutamatergic Synapse" enrichment within the “Cellular Component” GO term. **B.** Enrichment for the “Cellular Component” GO term within the “Glutamatergic Synapse" enrichment arranged by false discovery rate (FDR). Includes list of matching proteins in each network. **C.** Enrichment for the “Molecular Function” GO term within the “Glutamatergic Synapse" enrichment arranged by false discovery rate (FDR). Includes list of matching proteins in each network. **D.** Enrichment for the “Biological Process” GO term within the “Glutamatergic Synapse" enrichment arranged by false discovery rate (FDR). Includes list of matching proteins in each network. **E.** Enrichment for the Kyoto Genes and Genomes pathway within the “Glutamatergic Synapse" enrichment arranged by false discovery rate (FDR). Includes list of matching proteins in each network.

**Figure S1:** **Glutamatergic Synapse: NrbAP Pre and Post Synaptic Distribution** (**A**) Specific neurabin interacting proteins based on peptide spectral matches (PSMs) were input into the string database ([www.string-db.org](https://www.string-db.org/)). The proteins matching the “Glutamatergic Synapse” biological process term and their string interactions are plotted. Within this enrichment, proteins matching the “Presynapse”, “Postsynaptic Density”, and “Postsynapse”, were enriched. Proteins lacking enrichment are labeled “Misc.”. Network edges indicate confidence interaction score, and thicker edges indicate greater stronger evidence of association. Medium confidence (0.4) used as lower cutoff.

**Figure S2:** **Rotarod Performance in 8 Week Old Mice May Vary By Sex A**) Two way-RM ANOVA of male Nrb^+/+^ and Nrb^-/-^ mice demonstrates that both cohorts learn the accelerating rotarod task (F(4,24)= 21.22; p<0.0001) and male Nrb^+/+^ and Nrb ^-/-^ mice perform similarly to their littermate counterparts (F(1, 6)=1.719; p=0.2378) at the aggregate day level (**B**) Two way-RM ANOVA of male Nrb^+/+^ and Nrb^-/-^ demonstrates that both cohorts learn the accelerating rotarod task (F(2.947,17.68)= 9.376; p=0.0007) and Nrb^+/+^ and Nrb ^-/-^ perform similarly to their littermate counterparts (F(1, 6)=1.512; p=0.2648) at the individual trial level. (**C**) Two way-RM ANOVA of female Nrb^+/+^ and Nrb^-/-^ demonstrates that both cohorts learn the accelerating rotarod task (F(1.211, 7.267)= 7.175; p0.0270) but Nrb^-/-^ outperform their Nrb^+/+^ littermate counterparts (F(1, 6)=9.275; p=0.0226) at the aggregate day level. (**D**) Two way-RM ANOVA of female Nrb^+/+^ and Nrb^-/-^ demonstrates that both cohorts learn the accelerating rotarod task (F(3.105,18.63)= 4.860; p=0.0109) but Nrb^-/-^ trend towards performing better than their Nrb ^+/+^ littermate counterparts (F(1, 6)=3.987; p=0.0929) at the individual trial level. Mean ± SEM. Significant ANOVA results denoted by #p≤0.05, ##p≤0.01, ###p≤0.001, ####p≤0.0001.

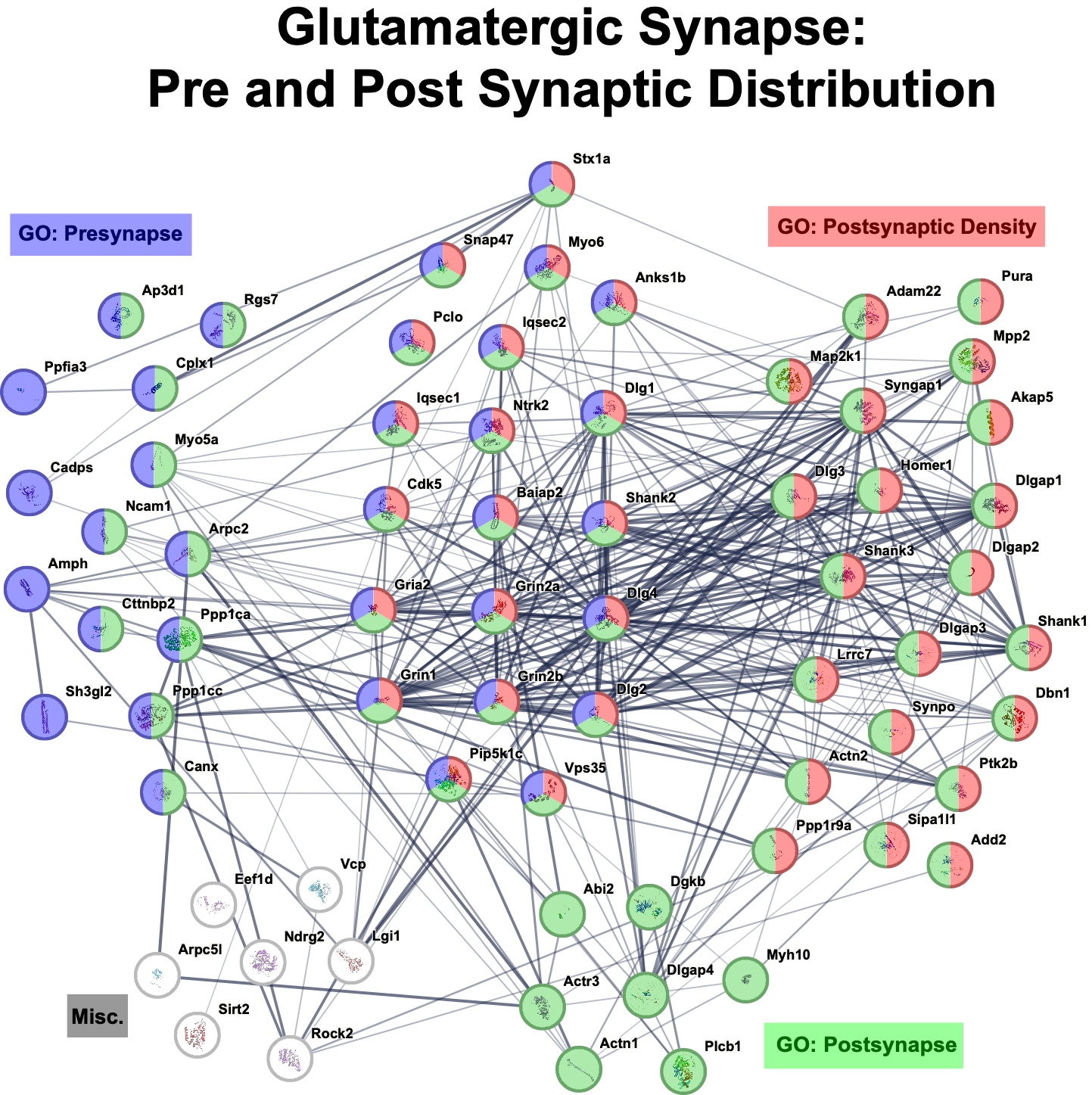
**Figure S1**

**Figure S2**

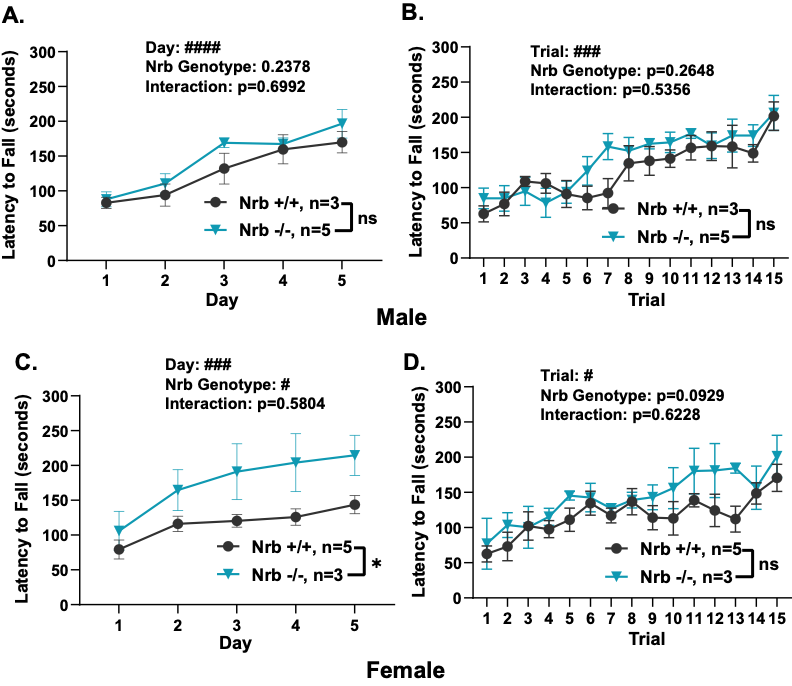
